# Rootstocks influence the response of ripening grape berries to leafroll associated viruses

**DOI:** 10.1101/2021.03.14.434319

**Authors:** Amanda M. Vondras, Larry Lerno, Mélanie Massonnet, Andrea Minio, Adib Rowhani, Dingren Liang, Jadran Garcia, Daniela Quiroz, Rosa Figueroa-Balderas, Deborah A. Golino, Susan E. Ebeler, Maher Al Rwahnih, Dario Cantu

## Abstract

Grapevine leafroll-associated virus (GLRaV) infections are accompanied by symptoms with varying severity. Using a dedicated experimental vineyard, we studied the responses to GLRaVs in ripening berries from Cabernet franc grapevines grafted to different rootstocks and with zero, one, or pairs of leafroll infection(s). RNA sequencing data were mapped to a high-quality Cabernet franc genome reference assembled to carry out this study and integrated with hormone and metabolite abundance data. This study identified several molecular levers that participate in responses to GLRaVs, including those that are condition-dependent. This included describing common responses to GLRaVs that were reproduced in two consecutive years, in plants grafted to different rootstocks, and in more than one infection condition. Though different infections were inconsistently distinguishable from one another overall, the effects of infections in plants grafted to different rootstocks were distinct at each developmental stage. Conserved responses included the modulation of pathogen detecting genes, increases in abscisic acid signaling and cytoskeleton remodeling gene expression. The abundance of abscisic acid (ABA), related metabolites, ABA and hormone signaling-related gene expression, and the expression of several transcription factor families differentiated rootstocks overall. These data show that rootstock influences the effect of GLRaVs in ripening berries.

## Introduction

Grapevine leafroll-associated viruses (GLRaVs) are among the most consequential pathogens affecting grapevine and have considerable economic impact (Atallah *et al*., 2012; Ricketts *et al*., 2015). GLRaVs are diverse and belong to the family *Closteroviridae,* with five different species and numerous strains in three genera (Naidu *et al*., 2015). Grapevines are often infected with several of these viruses simultaneously (Prosser *et al*., 2007; Rwahnih *et al*., 2009; Fuchs *et al*., 2021). Given their impact and global distribution, efforts to manage the spread of GLRaV, characterize their effects, and understand the interaction between the vine and pathogens have been undertaken.

Generally, plant responses to viruses include numerous changes in gene expression, gene regulation, and metabolism (Alazem & Lin, 2015; Moon & Park, 2016; Bester *et al*., 2016; Blanco-Ulate *et al*., 2017). Diverse pathogens and stresses elicit conserved responses from their hosts (Postnikova & Nemchinov, 2012; Shaik & Ramakrishna, 2013; Amrine *et al*., 2015; Jiang *et al*., 2015) and phylogenetically related viruses elicit similar responses (Rodrigo *et al*., 2012). Infections with GLRaVs have been associated with poorer fruit quality, lower yield, and leaves that curl, redden, and become brittle. Gene expression studies that implicate regulatory systems in the leafroll disease phenotype are few in number and have focused on the impact of GLRaV-3, highlighting changes in the expression of senescence-associated and flavonoid biosynthetic pathway genes (Espinoza *et al*., 2007b,a; Gutha *et al*., 2010; Vega *et al*., 2011). Additional transcriptomic study could help generate novel hypotheses concerning the controls that are fundamental to GLRaV responses (Mandadi & Scholthof, 2013; Gaiteri *et al*., 2013; Moon & Park, 2016); phytohormone signaling pathways and other molecular controls with far-reaching influence on the transcriptome are good candidates.

Though common responses might be expected in infected plants given the relatedness of GLRaVs, there is considerable variability in the severity of GLRaV infections. Some GLRaV infections appear asymptomatic or are mild (Kovacs *et al*., 2001; Poojari *et al*., 2013; Montero *et al*., 2016), but others cause significant changes in photosynthesis, metabolism, and gas exchange in leaves (Guidoni *et al*., 1997; 2000; Bertamini *et al*., 2004; Pereira *et al*., 2012; Endeshaw *et al*., 2014). Changes in fruit yield, organic and amino acids, titratable acidity, potassium, sugars, and flavonoids are also observed (Kliewer & Lider, 1976a; Cabaleiro *et al*., 1999; Lee & Martin, 2009; Lee *et al*., 2009; Alabi *et al*., 2016). These are influenced by host genotype (Kovacs *et al*., 2001; Montero *et al*., 2016), which virus or combination of viruses is present (Credi, 1997; Guidoni *et al*., 2000; Komar *et al*.), and environmental conditions (Cui *et al*., 2016).

Evidence relating GLRaV responses to rootstock is mixed (Golino, 1993; Komar *et al*., 2010). In a study of Cabernet franc vines grafted to different rootstocks, the effect of GLRaV infection on pruning weight depended on rootstock and the largest effects across all the parameters considered were observed in Kober 5BB-grafted vines (Rowhani, 2015). Similarly, fruit yield was influenced by both infection type and rootstock, with Kober 5BB-grafted vines most severely affected by a mixed infection with GLRaV-2, GLRaV-3, and *Grapevine fleck virus* (Golino *et al*., 2008b). In an extreme example, Red Globe scion buds infected with a strain of GLRaV-2 (formerly named *Grapevine rootstock stem lesion associated virus*) were used to inoculate Cabernet Sauvignon plants grafted to 18 different rootstocks; the infection was lethal in several rootstock conditions, including Kober 5BB (Uyemoto *et al*., 2001; Alkowni *et al*., 2011).

This study used Cabernet franc grapevines infected with zero, one, or two GLRaVs and grafted to two different rootstocks to identify responses to leafroll viruses in ripening berries that were conserved across experimental conditions and to determine whether or not GLRaV responses could be distinguished in the berries of plants grafted to different rootstocks. Grapevines were grown in a single experimental vineyard and berries were sampled during ripening in four consecutive years. Measures of vine growth and fruit composition were taken in the first two years. Total soluble solids (TSS) were measured in all four years. In the third and fourth years, RNA sequencing (RNAseq) and metabolite data were collected from Cabernet franc berries at four stages during ripening. RNAseq reads were mapped to the Cabernet franc genome, which was sequenced in long PacBio reads, assembled using the FALCON-Unzip pipeline, and scaffolded using Hi-C data. The same samples were used to measure stress and ripening-associated phytohormones, including abscisic acid (ABA), jasmonic acid (JA), and salicylic acid (SA), and ripening-related metabolites. Though many of the GLRaV effects occurred in individual years, a subset of reproducible conserved responses and rootstock-driven differences were discovered.

## Materials and methods

### Vineyard establishment

The experimental vineyard used in this study consists of Cabernet franc clone 04 (UC Davis, Foundation Plant Services, https://fps.ucdavis.edu/fgrdetails.cfm?varietyid=355; accessed March 2, 2021) grapevines grafted on different rootstocks and infected with zero, individual, or pairs of GLRaVs. All rootstocks and Cabernet franc scions were tested for grapevine pathogens. Total nucleic acid (TNA) extracts were prepared from all rootstocks and Cabernet franc scions as described by Al Rwahnih *et al*. (2017) (Rwahnih *et al*., 2017). Briefly, approximately 0.2 grams of leaf petioles were homogenized using a HOMEX 6 homogenizer (Bioreba AG, Reinach, Switzerland) and TNA extracts were prepared using a MagMAX-96 viral RNA isolation kit (ThermoFisher Scientific) per the manufacturer’s protocol. Extracted TNA samples were analyzed by reverse transcription quantitative PCR (RT-qPCR) using TaqMan probes on the QuantStudio 6 Flex Real-Time PCR System (ThermoFisher Scientific) as described previously (Osman *et al*., 2008; Klaassen *et al*., 2011; Rwahnih *et al*., 2017). The samples were screened for the following pathogens: *Grapevine leafroll-associated virus-1*, *Grapevine leafroll-associated virus-3*, *Grapevine leafroll-associated virus-4* (plus strains 5, 6, 9, Pr and Car; genus *Ampelovirus*), *Grapevine leafroll-associated virus-2* (plus strain 2RG; genus *Closterovirus*), *Grapevine leafroll-associated virus-7* (genus *Velarivirus*), *Grapevine fleck virus* (genus *Maculavirus*), *Grapevine rupestris vein feathering virus* (genus *Marafivirus*), *Grapevine fanleaf virus*, *Tobacco ringspot virus*, *Tomato ringspot virus* (genus *Nepovirus*), *Grapevine virus A*, *Grapevine virus B*, *Grapevine virus D*, *Grapevine virus E*, *Grapevine virus F*, (genus *Vitivirus*), *Grapevine red blotch virus* (genus *Grablovirus*), *Grapevine rupestris stem-pitting associated virus* (genus *Foveavirus*), phytoplasmas and *Xylella fastidiosa*, the causative agent of Pierce’s disease.

In Fall 2008, Cabernet franc grapevines were bench-grafted onto rootstocks, including Millardet et de Grasset (MGT) 101-14 and Kober 5BB. These plants were subsequently grown in a greenhouse. Between 2009 and 2011, the rootstock portions of these plants were inoculated with two chip buds from single leafroll-infected plants. Infected plants used for chip buds were reconfirmed by RT-qPCR. Plants infected with a single species of GLRaV received two identical chip buds. Plants infected with two species of GLRaVs also received two chip buds, each carrying a single virus. Plants infected with GLRaV-1, GLRaV-2 and/or GLRaV-3 were inoculated with two or more isolates of each species of GLRaV. The inoculated plants were kept in a greenhouse for approximately one month, acclimatized, then were planted in the field. Healthy controls included non-chip budded plants and plants chip budded from a healthy source.

The vines were planted according to a randomized complete block design, with seven feet between vines and nine feet between rows. One group of five vines was planted per rootstock × infection condition in each of three blocks. Healthy vines were distributed throughout each block to monitor the spread of viruses, and experimental vines were sampled yearly to test and reaffirm the vines’ infection status. A buffer zone of healthy vines was planted as a barrier between the leafroll vineyard and other vineyards in the area. Vines were trained with a bilateral cordon and spur pruned.

### Cabernet franc genome sequencing and assembly

High quality genomic DNA were isolated from grape leaves using the method described in Chin *et al*. (2016). DNA purity was evaluated with a Nanodrop 2000 spectrophotometer (Thermo Scientific, Hanover Park, IL), quantity with a Qubit 2.0 Fluorometer (Life Technologies, Carlsbad, CA), and integrity by electrophoresis. SMRTbell libraries were prepared for Cabernet franc clone 04 as described by Massonnet *et al*. (2020). Final libraries were evaluated for quantity and quality using a Bioanalyzer 2100 (Agilent Technologies, CA) and sequenced on a PacBio RS II (DNA Technology Core Facility, University of California, Davis).

*De novo* assembly of Cabernet franc clone 04 (UC Davis, Foundation Plant Services, https://fps.ucdavis.edu/fgrdetails.cfm?varietyid=355; accessed March 2, 2021) was performed using PacBio RS II data and FALCON-Unzip (v. 1.7.7; (Chin *et al*., 2016). Repetitive sequences were masked before and after read error-correction using TANmask and REPmask modules in Damasker (Myers, 2014). Multiple FALCON-Unzip parameters were tested to optimize the assembly as described in Minio *et al.,* 2019(Minio *et al*., 2019a). Haplotype reconstruction was performed with default parameters. Finally, contigs were polished with Quiver (Pacific Biosciences, bundled with FALCON-Unzip v. 1.7.7).

The primary assembly was scaffolded to reduce sequence fragmentation. First, primary contigs were scaffolded with SSPACE-LongRead (v.1.1; (Boetzer & Pirovano, 2014). Junctions supported by at least 20 reads (“-l 20”) were allowed. Hi-C data and the proprietary HiRise software (v1.3.0-1233267a1cde) were used for hybrid scaffolding. A Dovetail Hi-C library was prepared by Dovetail Genomics (Scotts Valley, CA, USA) as described in Lieberman-Aiden *et al*. (2009) (Lieberman-Aiden *et al*., 2009) and sequenced on an Illumina platform, generating 2 × 150bp paired-end reads.

The repeat and gene annotation were performed as reported in Vondras *et al*. (2019) (Vondras *et al*., 2019). Briefly, RepeatMasker (v. open-4.0.6; (Smit *et al*.)) and a custom *V. vinifera* repeat library (Minio *et al*., 2019b) were applied to identify repetitive elements in the genome. To annotate genes, publicly available datasets were used as evidence for gene prediction. Transcriptional evidence included *Vitis* ESTs, Cabernet Sauvignon corrected Iso-Seq reads (Minio *et al*., 2019b), Tannat (Da Silva *et al*., 2013), Corvina (Venturini *et al*., 2013), and Cabernet Sauvignon transcriptomes (Massonnet *et al*., 2020), and previously published RNAseq data (PRJNA260535). Swissprot viridiplantae data and *Vitis* data were used as experimental evidence. Each RNAseq sample was trimmed with Trimmomatic (v. 0.36; (Bolger *et al*., 2014)) and assembled with Stringtie (v. 1.3.3; (Pertea *et al*., 2015)). This data was mapped on the genome using Exonerate (v. 2.2.0, transcripts and proteins; (Slater & Birney, 2005)) and PASA (v. 2.1.0, transcripts; (Haas *et al*., 2003)). Alignments and *ab initio* predictions generated with SNAP (ver. 2006-07-28; (Korf, 2004)), Augustus (ver. 3.0.3; (Stanke *et al*., 2006)), and GeneMark-ES (ver. 4.32; (Lomsadze *et al*., 2005)) were used as input for EVidenceModeler (v. 1.1.1; (Haas *et al*., 2008)). EVidenceModeler was used to identify consensus gene structures. A functional annotation was obtained using the RefSeq plant protein database (ftp://ftp.ncbi.nlm.nih.gov/refseq, retrieved January 17th, 2017; (Jones *et al*., 2014)) as in Minio *et al*. (2019).

### Sampling and sample preparation

Berries from Cabernet franc grapevines grafted to Kober 5BB or MGT 101-14 and infected with either GLRaV-1 (GLRaV-1 (+)), GLRaV-3 (GLRaV-3 (+)), GLRaV-4 (strain 5; GLRaV-4 (+)), GLRaV-1 and GLRaV-2 (GLRaV-1,2 (+)), or GLRaV-1 and GLRaV-3 (GLRaV-1,3 (+)) were sampled during ripening in 2017 and 2018. In both years, fruits were sampled at pre-véraison, véraison, mid-ripening, and at commercial harvest. These stages correspond to modified Eichhorn-Lorenz (Coombe, 1995) stages 34 (green berries beginning to soften and °Brix starts increasing), 35 (berries begin to change color and enlarge, ∼50% of berries within clusters show color transition), 36/37 (berries with intermediate °Brix values / berries are not quite ripe), and 38 (berries are harvest-ripe). Effort was made to sample fruits in 2018 at developmental stages comparable to 2017, as determined by TSS. In 2017, berries were sampled July 7^th^, July 31^st^, August 14^th^, and August 31^st^. In 2018, berries were sampled June 28^th^, July 30^th^, August 13^th^, and August 30^th^.

On each sampling date, six biological replicates were taken, with berries from one plant constituting one biological replicate. Two biological replicates per condition were sampled from each of three blocks. There were two exceptions. For Kober 5BB-grafted GLRaV-1,2 (+), three biological replicates were drawn from each of two plants in one block. For Kober 5BB-grafted GLRaV-3 (+), two of the six biological replicates were drawn from one plant. Approximately twenty berries were sampled per plant, with equal numbers of berries sampled from each side of the vine. Samples were then temporarily cooled on ice.

Samples were processed immediately following sampling. This included rinsing berries with deionized water, deseeding them, and snap-freezing them in liquid nitrogen. Berries were stored at −80 °C until crushed into a fine powder while frozen using a mechanical mill. Total soluble solids were subsequently measured in technical triplicate using a digital refractometer. Four of six biological replicates, a total of 192 samples per year, were used for subsequent RNA sequencing and LC-MS.

### RNA extraction and sequencing library preparation

Total RNA was extracted from two grams of finely ground berry pericarp tissue as previously described (Blanco-Ulate *et al*., 2013). RNA purity was evaluated with a Nanodrop 2000 spectrophotometer (Thermo Scientific, Hanover Park, IL), quantity with a Qubit 2.0 Fluorometer (Life Technologies, Carlsbad, CA), and integrity by electrophoresis. RNAseq libraries were prepared using the Illumina TruSeq RNA sample preparation kit v.2 (Illumina, CA, USA) and barcoded individually following the manufacturer’s protocol. Final libraries were evaluated for quantity and quality with the High Sensitivity chip in a Bioanalyzer 2100 (Agilent Technologies, CA).

Libraries were sequenced as 100 base-pair, single-end reads, using an Illumina HiSeq4000 sequencer (DNA Technology Core Facility, University of California, Davis) producing an average of 18.07M ± 4.56M reads per sample in 2017 and 14.69M ± 2.11M reads per sample in 2018.

### RNAseq data analysis

RNAseq reads were trimmed using Trimmomatic (v. 0.36; (Bolger *et al*., 2014)) and the following settings: Leading:2, Trailing:2, Sliding window:4:20, Min length:70. Reads were mapped to the primary assembly of the Cabernet franc genome using HISAT2 (-k 1; v. 2.0.5; (Kim *et al*., 2015)) and counts were generated using htseq-count with default parameters (v. 0.9.0; (Anders *et al*., 2015)).

All subsequent analyses were done using R (R Core Team, 2020) in the R Studio environment (R Studio Team, 2020). Data normalization and differential expression analyses were done using DESeq2 (v. 1.24.0; (Love *et al*., 2014)). A variance stabilizing transformation (VST) was applied to expression data using DESeq2. VST data were centered in each GLRaV (+) condition relative to the mean expression per gene in GLRaV (-) given the same time, rootstock, and year.

Centered data were used for Multiple Factor Analysis (MFA). The FactoMineR R package was used for MFA (Le *et al.,* 2008). Genes were included in the MFA if they were differentially expressed (1) versus GLRaV (-) and/or if the effects of an infection differed between rootstocks (2) in at least one year (P < 0.05), and (3) if the direction of the effect relative to GLRaV (-) was consistent in both years, even if a significant effect was only observed in one year. An MFA was repeated for each developmental stage separately. All hormone and metabolite data were included.

Statistical overrepresentation tests were done using the clusterProfiler R package (Yu *et al.,* 2012) and VitisNet functional categories (Grimplet *et al*., 2009). To make use of the VitisNet functional annotations, Cabernet franc genes were used to query Pinot noir PN40024 sequences with blastp (Altschul *et al*., 1990). The best hits, with no less than 80% reciprocal identity and coverage, were retained.

A curated list of ABA biosynthetic pathway and signaling genes annotated in PN40024 was retrieved from Pilati *et al.,* 2017 (Pilati *et al*., 2017)(Supplementary File S1). As described above, Cabernet franc genes were used to query Pinot noir PN40024 sequences with blastp (Altschul *et al*., 1990). The best hits were retained; all target-query pairs had no less than reciprocal 97% identity and 86% reciprocal coverage (median coverage, 99%).

### Hormone extraction and LC-MS/MS

Approximately fifty milligrams (mean 50.86 ± 3.33 mg) of berry powder were weighed for the extraction and quantitation of ABA, SA, and JA. Exact weights were recorded to later calculate the exact amount of each of these analytes per milligram of fresh tissue. The same samples used for RNA sequencing were used for this analysis. Four biological replicates were used. Extractions and analyses were randomized and performed in technical duplicate.

Hormones were extracted following a method described by Pan *et al.,* (2010) with a few modifications (Pan *et al*., 2010). Samples were subjected to 500 mL of 2-propanol:H_2_O: HCl (2:1:0.002) and spiked with 50 ng of d_6_ abscisic acid, d_5_ jasmonic acid, and d_4_ salicylic acid (CDN Isotopes, Pointe-Claire, Quebec, Canada). Samples were vortexed then placed in an ultrasonic ice bath for 30 minutes, then washed with dichloromethane and centrifuged at 13,200 rpm for 5 minutes. Nine-hundred microliters of the lower phase were taken and dried. Samples were reconstituted in 100 microliters of 15% methanol and stored at −20 °C until analysis.

Chromatographic separations were performed on an Agilent 1290 Infinity UHPLC with binary pump, autosampler, and temperature-controlled column compartment. Samples were analyzed in multiple reaction mode using an Agilent (Santa Clara, CA, USA) 6460 triple quadrupole mass spectrometer equipped with an Agilent Jetstream electrospray source. Instrument control and data acquisition was performed in Agilent Masshunter Acquisition (ver 7). Hormones were detected using the MRM transitions in Supplementary File S2. Hormones were quantified by internal standard after determining the linear regions for each.

A binary solvent system consisting of 10 mM ammonium formate in water (mobile phase A) and 10 mM ammonium formate in acetonitrile (mobile phase B) was used for chromatographic separations on an Agilent (Santa Clara, CA, USA) Poroshell 120 EC-C18 (150 × 2.1 mm, 2.7 µm) column fitted with a Poroshell guard column containing the same phase. The mobile phase program used for separations was as follows: 0 min, 5% B; 2 min, 5% B; 6 min, 40% B; 6.5 min, 100% B; 8.5 min, 100% B; 9.5 min, 5% B. The column was allowed to equilibrate for 6 minutes at starting conditions prior to the next injection. Flowrate for analyses was 0.4 mL/min, the column temperature was maintained at 35 °C, and an injection volume of 20 µL was used for all samples. Source conditions for the Jetstream electrospray source were as follows: Drying gas temperature 100 °C, drying gas flow 8 L/min, sheath gas temperature 300 °C, sheath gas flow 10 L/min, nebulizer 50 psi, capillary voltage 3500 V (both positive and negative mode), and nozzle voltage 0 V.

### Water-soluble metabolite extraction and LC-MS

The same samples used for RNAseq and hormone analyses were used to measure water-soluble metabolites by LC-MS. Approximately two hundred milligrams (mean 202.12 ± 4.87 mg), of berry powder were weighed while frozen. Extractions and analyses were randomized and performed in technical duplicate. Hormones were extracted in 1 mL 1% HCl in HPLC-grade H_2_O (prepared in-house, final resistance 18 MΩ, 0.2 µm filtered) and spiked with salicin (Sigma Aldrich, St. Louis, MO, USA) as an internal standard (50 µg/L final concentration). The samples were vortexed, placed in an ultrasonic ice bath for 30 minutes, and centrifuged. The supernatants were collected and frozen at −80°C until analyzed.

Chromatographic separations were performed on an Agilent (Santa Clara, CA, USA) 1290 Infinity II UHPLC (equipped with a binary pump, multisampler, and temperature-controlled column compartment) that was connected to a 6545 Q-TOF MS. An Agilent 1260 isocratic pump was used for introduction of reference mass solution (constant flow of 0.8 mL/min during analysis and split 1:100 prior to reference nebulizer). Chromatographic separations were performed using an Agilent Zorbax Eclipse Plus C18 (50×2.1 mm, 1.8 µm) column fitted with a guard column of the same stationary phase and kept at 50 °C. All extracts were analyzed on an Agilent (Santa Clara, CA, USA) 6545 Q-TOF mass spectrometer equipped with a dual Jetstream electrospray source, once in positive mode and once in negative mode. Instrument control and data acquisition was performed in Agilent Masshunter Acquisition (ver 8). Prior to analysis, a low mass transmission tune was performed in both positive and negative ion modes following the Agilent guidelines. Data acquisition was performed in the MS only mode with a mass spectral range of *m/z* 90-1000 and a scan rate of 4 spectra/sec. The Agilent All-ions workflow with collision energies of 0 V and 25 V was used to collect non-targeted fragmentation data for compound identification.

A binary solvent system was used consisting of 1% formic acid in water (mobile phase A) and 1% formic acid in acetonitrile (mobile phase B). The solvent program was as follows: 0.00 min, 3% B; 4.67 min, 35% B; 5.00 min, 80% B; 6.67 min, 80% B; 7 min, 3% B. The column was equilibrated at starting conditions for 3 minutes between injections and an injection volume of 4 µL was used for all samples. Reference mass correction was performed using purine and hexakis (1H,1H,3H-tetrafluoropropocy)phosphazene for both positive and negative modes. Water soluble metabolites (Supplementary File S2) were analyzed by a targeted data analysis method in Masshunter Quantitative Analysis (ver 10.1). Metabolites were identified by accurate mass, retention time, and characteristic fragments from the 25 V collision energy channel.

### Data visualization

Figures depicting quantitative data were built using the following R packages: ggplot2 (Wickham, 2016), hrbrthemes (Rudis, 2020), wesanderson (Ram and Wickham, 2018), pheatmap (Kolde, 2019) and UpSetR (Gehlenborg, 2019).

## Results

### GLRaV species and rootstock influence canopy density, cluster weight and fruit composition

The ways that individual and pairs of GLRaVs (GLRaV-1, GLRaV-3, GLRaV-4 strain 5, GLRaV-1,2, and GLRaV-1,3) and rootstock (MGT 101-14 and Kober 5BB) combinations influence grape berry ripening were studied in a dedicated experimental vineyard at the University of California, Davis.

Typical grapevine leafroll disease symptoms (i.e., leaf-reddening and curling) were observed by mid-ripening (Fig. 1A). In addition, there was a visible, stark reduction in canopy density and cluster size in GLRaV-1,2 (+) versus GLRaV (-) in vines grafted to Kober 5BB that was not readily apparent in vines grafted to MGT 101-14 with the same infection status (Fig. 1B).

**Figure 1.**
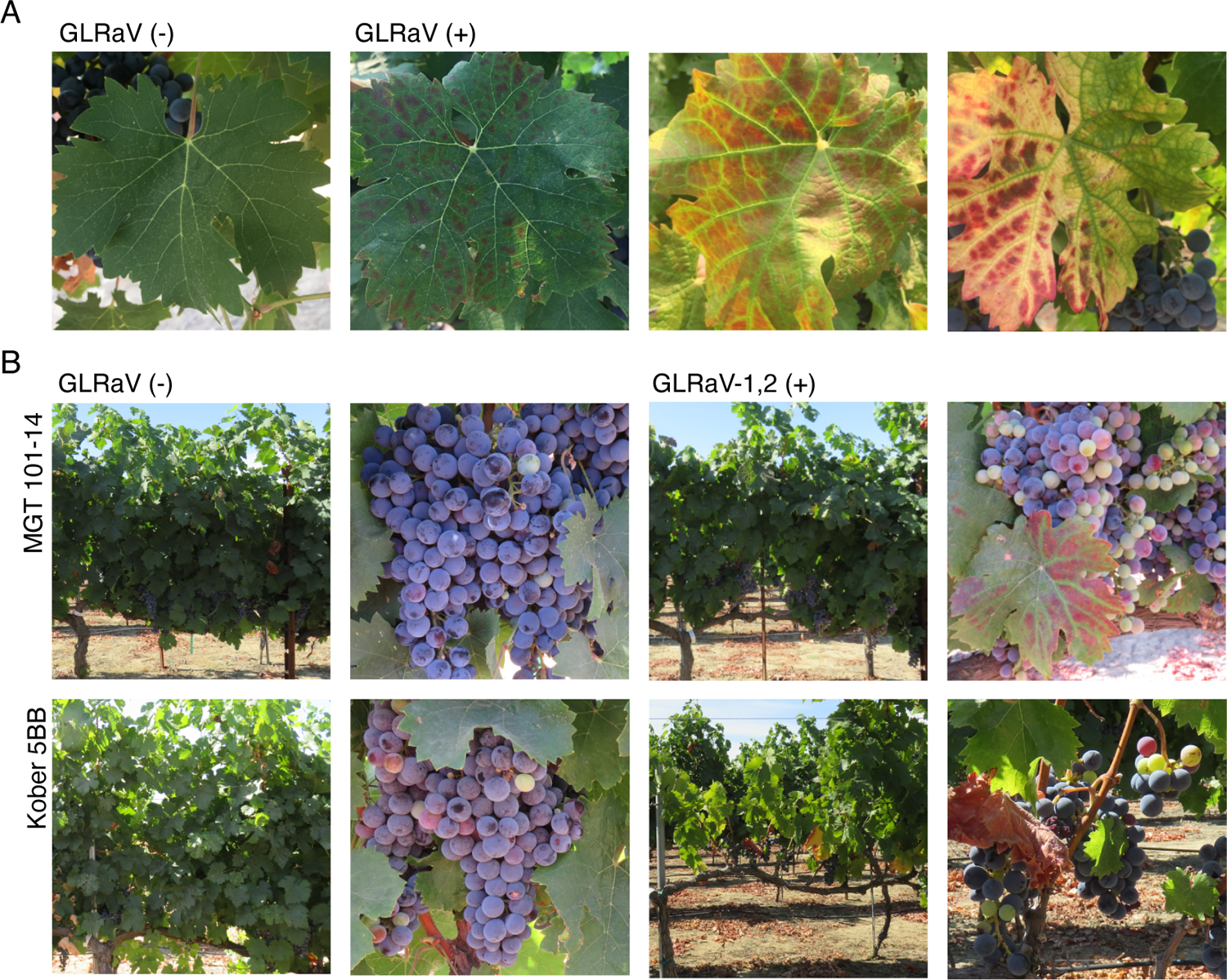
Effects of grapevine leafroll-associated viruses on Cabernet franc leaves, canopy density, and cluster size. **(A)** Leaves sampled slightly before (August 8) and at mid-ripening (August 13) in 2018 from GLRaV (-) and GLRaV (+) **(B)** Canopy and berry clusters from GLRaV (-) and GLRaV-1,2 (+) in different rootstock conditions on August 14, 2017.

Vine growth, cluster weight, and other measures were collected in 2015 and 2016 (Fig. 2, Supplementary Fig. S3). The effect of GLRaV infection on dormant pruning weight, berry weight, pH, and tartaric acid in 2015 and on moisture content, total anthocyanin content, and titratable acidity in 2016 differed significantly based on the rootstock present (ANOVA, P < 0.05). This interaction was significant for malic acid in 2015 and 2016. Significant differences in dormant pruning weights, total cluster weights, and tartaric acid were observed in plants with different GLRaV infection status and rootstock (Tukey HSD, P < 0.05). In contrast, few or no significant differences between GLRaV (+) given the same rootstock were observed for total anthocyanins, moisture content, malic acid, pH, titratable acidity, weight per berry, or yeast assimilable nitrogen and overwhelmingly in a single year if at all (Supplementary Fig. S3).

**Figure 2.**
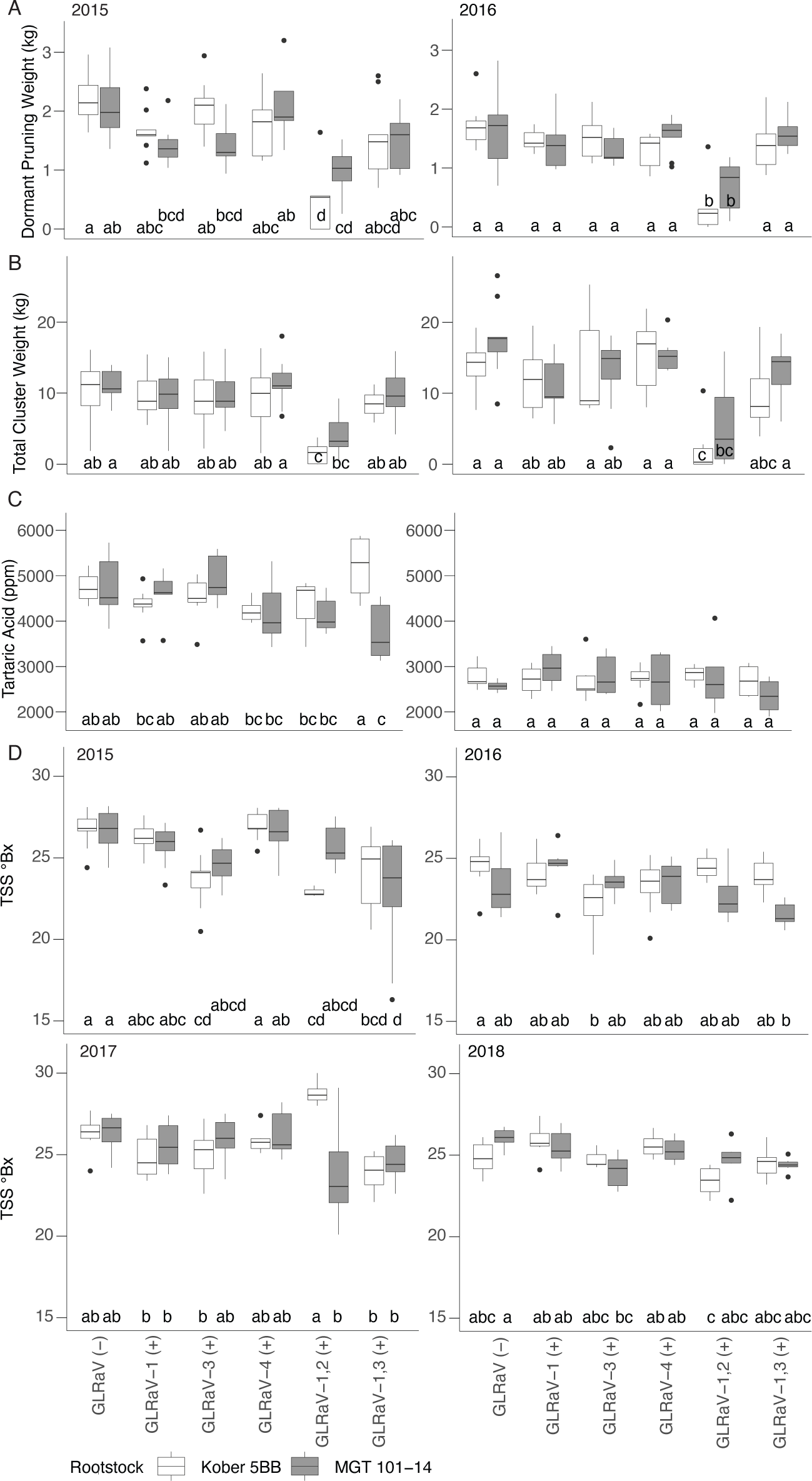
Effects of GLRaV infection on **(A)** Dormant Pruning Weight, 5 ≤ n ≤ 9, **(B)** Total cluster weight, 5 ≤ n ≤ 9 and **(C)** Tartaric acid, 3 ≤ n ≤ 9, in 2015 and 2016. No significant differences between groups were observed in tartaric acid in 2016. **(D)** Total soluble solids at harvest in four consecutive years. 2015-2016, 5 ≤ n ≤ 9; 2017-2018, n = 6. Group differences are indicated with non-overlapping letters (Tukey HSD, P < 0.05).

Overall, GLRaV infection tended to reduce dormant pruning weight and cluster weight. The dormant pruning weights and cluster weights of GLRaV-1,2 (+) were significantly less than GLRaV (-) and other GLRaV (+) (Fig. 2AB); this was observed for both rootstock conditions. Significant differences in fruit tartaric acid levels were observed in only 2015 and were between GLRaV-1,3 (+) grafted to different rootstocks and between plants with different GLRaV infection status (Fig. 2C). In each year except 2015, there was a significant interaction between rootstock and GLRaV infection status in terms of TSS at harvest (ANOVA, P < 0.05, Figure 2D). This interaction was significant at each other developmental stage in 2017 and at pre-véraison in 2018 (Supplementary Fig. S3). Significant differences in TSS at harvest were found between GLRaV (-) and GLRaV (+) in each year except 2017 (Fig. 2D). Overall, significant reductions in TSS relative to GLRaV (-) were limited to the dual infections and GLRaV-3 (+). Significant differences were observed between rootstocks in GLRaV 1,2 (+) at every developmental stage, albeit only in 2017 (Fig. 2D, Supplementary Fig. S3).

These data provide limited evidence that (1) different GLRaVs may or may not affect various aspects of vine growth and fruit composition, (2) that some of these differences are rootstock-specific, and (3) that although some of these effects are observed across years, year-to-year differences may impact whether or not effects occur.

### GLRaVs elicit reproducible changes in gene expression across infection types and rootstocks

We used RNAseq to sequence the transcriptome of 384 Cabernet franc berry samples collected from plants grafted to different rootstocks (Kober 5BB or MGT 101-14), with different GLRaV infection status (GLRaV (-), GLRaV-1 (+), GLRaV-3 (+), GLRaV-4 (+), GLRaV-1,2 (+) or GLRaV-1,3 (+)), at four developmental stages (pre-véraison, véraison, mid-ripening, and harvest), and in two consecutive years (2017 and 2018).

Because of the remarkable structural and gene content variability among grape cultivars (Da Silva *et al.,* 2013; Venturini *et al.,* 2013; Minio *et al.,* 2019), we built a genome reference specifically for the analysis of these RNAseq data. The genome was assembled into 504 primary contigs (N50 = 5.74 Mbp) for a total assembly size of 570 Mbp. This is comparable to the size of the Zinfandel (591 Mbp; (Vondras *et al*., 2019)), Cabernet Sauvignon (590 Mbp; (Chin *et al*., 2016)), Chardonnay (490 Mb; (Roach *et al*., 2018)) and Pinot noir PN40024 (487 Mb; (Jaillon *et al*., 2007)) genomes. Three thousand eighty-five additional haplotigs were assembled with an N50 of 184 kbp (Supplementary Table S4). The primary assembly and haplotigs were annotated with 33,563 and 19,146 protein-coding genes, respectively (Supplementary Table S4).

Ripening was associated with transcriptomically distinct developmental stages. Samples clustered primarily by developmental stage and secondarily by year, though samples at harvest clustered separately (Supplementary Fig. S5). Genes with comparable, significant responses (consistently up-regulated or down-regulated, P < 0.05) in both years of the study were selected to identify reproducible responses to GLRaVs during ripening. Gene expression in GLRaV (+) was compared to gene expression in GLRaV (-) grafted to the same rootstock at the same developmental stage (Fig. 3A). In addition, the effects of each GLRaV infection on gene expression at each developmental stage were compared in plants grafted to different rootstocks (Supplementary Fig. S6).

**Figure 3.**
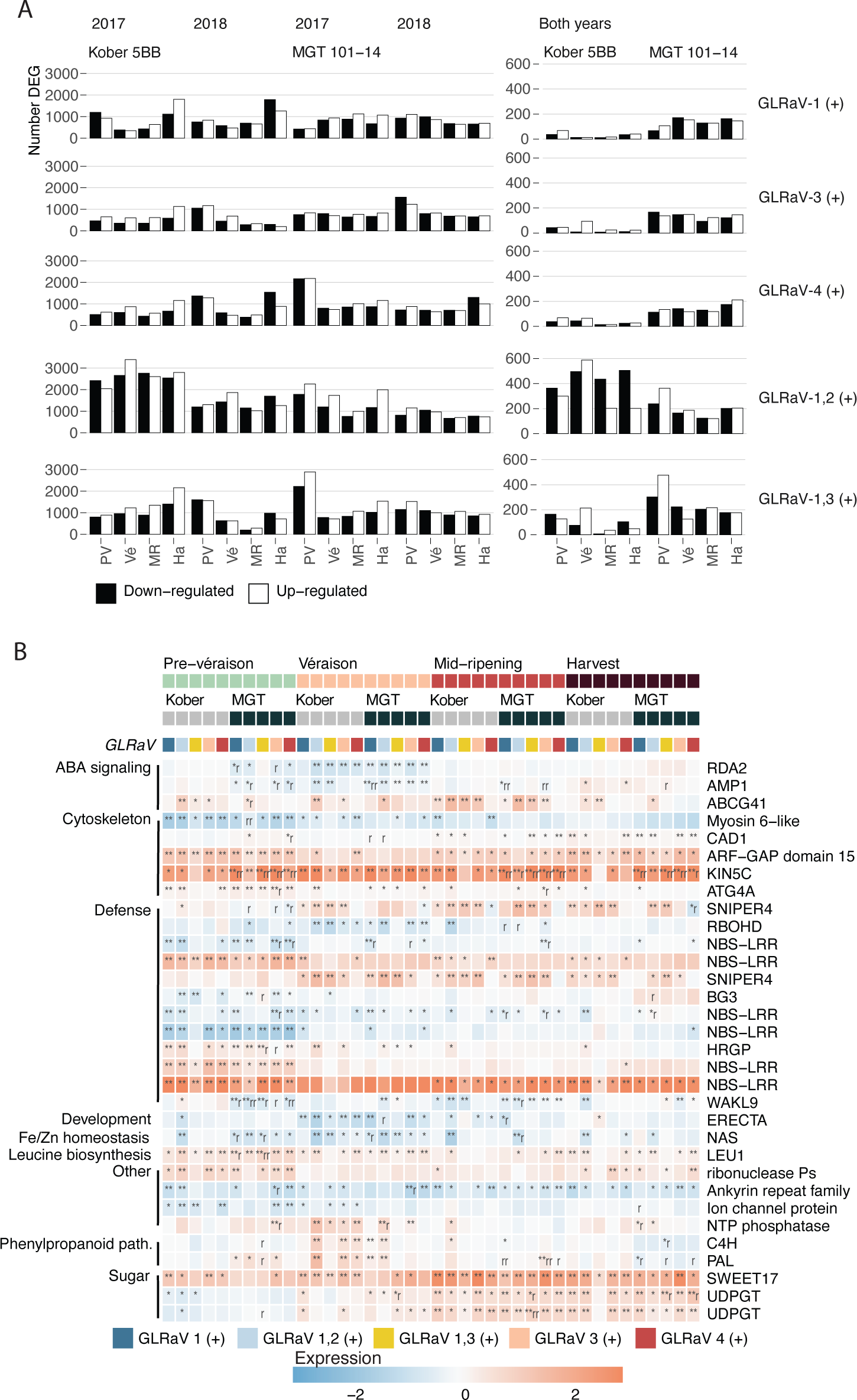
Differentially expressed genes in GLRaV-infected grapevines. **(A)** Barplots showing the number of differentially expressed (P < 0.05) genes up and down-regulated in GLRaV (+) versus GLRaV (-) in berries from Cabernet franc grapevines grafted to different rootstocks (Kober 5BB, Kober; MGT 101-14, MGT) at each developmental stage (Pre-véraison, PV; Véraison, Vé; Mid-ripening, MR; Harvest, Ha) in each year and in both years. **(B)** Conserved responses to GLRaVs (P < 0.05) observed in both rootstock conditions and in more than one GLRaV (+) condition. One asterisk, differentially expressed in one year; Two asterisks, differentially expressed in both years. One or two “r”, the effects of a particular GLRaV infection differ between rootstocks at the same developmental stage. Any notation requires the direction (up/down-regulation) of the effect to be consistent in both years, even if a significant change occurred in only a single year.

On average, 7.1% of the genes differentially expressed between GLRaV (-) and GLRaV (+) were reproduced in both years (Fig. 3A). This percentage was slightly above average for plants with dual, relatively more severe, infections (8.6%, GLRaV-1,3 (+); 8.9%, GLRaV-1,2 (+)) and below average for individual infections (5.8%, GLRaV-1 (+); 6% GLRaV-4 (+); 6.2% GLRaV-3 (+)). A subset of 32 genes significantly changed their expression level in two or more GLRaV(+) infection conditions, in both rootstock conditions, and at least one developmental stage (Fig. 3B). These genes constitute the “conserved” responses to GLRaVs in Cabernet franc berries during ripening.

The majority of these are associated with defense, ABA signaling, and cytoskeleton organization and biogenesis (Fig. 3B; Supplementary File S7). Six of these were nucleotide-binding site and leucine-rich repeat-containing (NBS-LRR) genes; half of these were up-regulated. Two F-box genes encoding SNIPER4 were up-regulated (Huang *et al*., 2018), as was a gene encoding a hydroxyproline-rich glycoprotein (HRGP). HRGP and NBS-LRR proteins are associated with pathogen detection (DeYoung & Innes, 2006). Genes encoding a respiratory burst oxidase protein D (RBOHD), a wall-associated kinase-like protein (WAKL), and a beta-glucosidase 3 (BG3) were down-regulated. RBOHD participates in the production of reactive oxygen species (ROS) and hypersensitive responses to pathogens (Otulak-Kozieł *et al*., 2019). RBOH family proteins are targeted by Snf1-related kinase 2 (SnRK2) phosphorylation, a key component of the ABA signaling pathway. Likewise, a *WAKL* gene in citrus participates in jasmonic acid and ROS signaling (Li *et al*., 2020). Among the functions of BGs are the activation of ABA and SA by freeing them from the conjugates that render them inactive (Seo *et al*., 1995; Morant *et al*., 2008; Zhang *et al*., 2012; Sun *et al*., 2014; Jia *et al*., 2016). Several ABA-related genes were among the conserved GLRaV responses, including an up-regulated ABC transporter and two down-regulated genes, *AMP1* and *RDA2*. *AMP1* negatively regulates ABA sensitivity (Shi *et al*., 2013). *RDA2* participates in the inhibition of ABA signaling and the promotion of MAPK signaling (Park *et al*., 2019).

There were five genes related to the cytoskeleton sensitive to GLRaV infection. Only one of these, a myosin VI motor protein coding gene, was down-regulated. The four others were autophagy genes (*ATG) 4A* and *constitutively activated cell death 1* (*CAD1)*, which function in autophagy, lytic pore formation, and hypersensitive response (Yoshimoto *et al*., 2004; Morita-Yamamuro *et al*., 2005; Haxim *et al*.), *Kinesin-like 5C (KIN5C)*, which encodes a microtubule motor protein (Reddy & Day, 2001), and an ADP-ribosylation factor GTPase-activating protein (*ARF-GAP*) domain 15, which helps efficiently load vesicles and remodel the actin cytoskeleton (Inoue & Randazzo, 2007).

Several additional general functional categories were present among the 32 genes that exhibited conserved responses to GLRaVs (Fig. 3B). Genes encoding phenylalanine ammonia-lyase (PAL) and cinnamate 4-hydroxylase (C4H), which catalyze the first two steps of the phenylpropanoid pathway, two genes encoding UDP glucosyltransferases (UDPGT), which conjugate sugars, and *SWEET17*, a sugar transporter, were up-regulated, as was a gene encoding 3-isopropylmalate dehydratase, an enzyme in the leucine biosynthetic pathway. Two genes, encoding a leucine-rich repeat receptor-like kinase (LRR-RLK) called ERECTA and nicotianamine synthase (NAS), were down-regulated. *ERECTA* participates in organ development and resistance to bacterial and fungal pathogens (Goff & Ramonell, 2007). *NAS* expression increases Fe and Zn abundance in rice (Moreno-Moyano *et al*., 2016; Nozoye, 2018).

Generally, these genes and their changes in expression suggest that a common response to GLRaVs in Cabernet franc berries during ripening includes the modulation of pathogen detecting genes, an increase in ABA transport and signaling, a decrease in ROS-related signaling, and an enhancement of cytoskeleton remodeling, vesicle trafficking, phenylpropanoid metabolism, sugar transport and conjugation, and leucine biosynthesis.

### Rootstock influences the impact of GLRaV on abscisic acid metabolism

The same berry samples used for RNAseq were used to measure three hormones associated with ripening and/or stress, including SA, JA, and ABA, and additional metabolites, including xanthoxin, a precursor to ABA, and ABA glucose ester (ABA-GE), a conjugate of ABA implicated in its long-distance transport (Jiang & Hartung, 2008). The mean levels of SA and JA were significantly influenced by year and/or by interactions between year, rootstock, and GLRaV at pre-véraison (ANOVA, P < 0.05), but no significant differences were observed between individual groups (Tukey HSD, P > 0.05) (Supplementary File S8). In contrast, year alone had a significant impact on the levels of ABA and related metabolites measured at each developmental stage, but largely did not interact with rootstock or GLRaV infection type to affect the abundance of ABA and related metabolites (Supplementary File S8). In addition, the effect of GLRaV infection on ABA and ABA-GE content significantly differed based on rootstock (Supplementary File S8). Significant differences in ABA and ABA-GE content were observed between rootstocks in plants with identical infection status and between plants with different GLRaV status grafted to the same rootstock (Tukey HSD, P < 0.05). Such differences were scarcely observed for xanthoxin, a precursor to ABA (Fig. 4A, Supplementary File S8).

**Figure 4.**
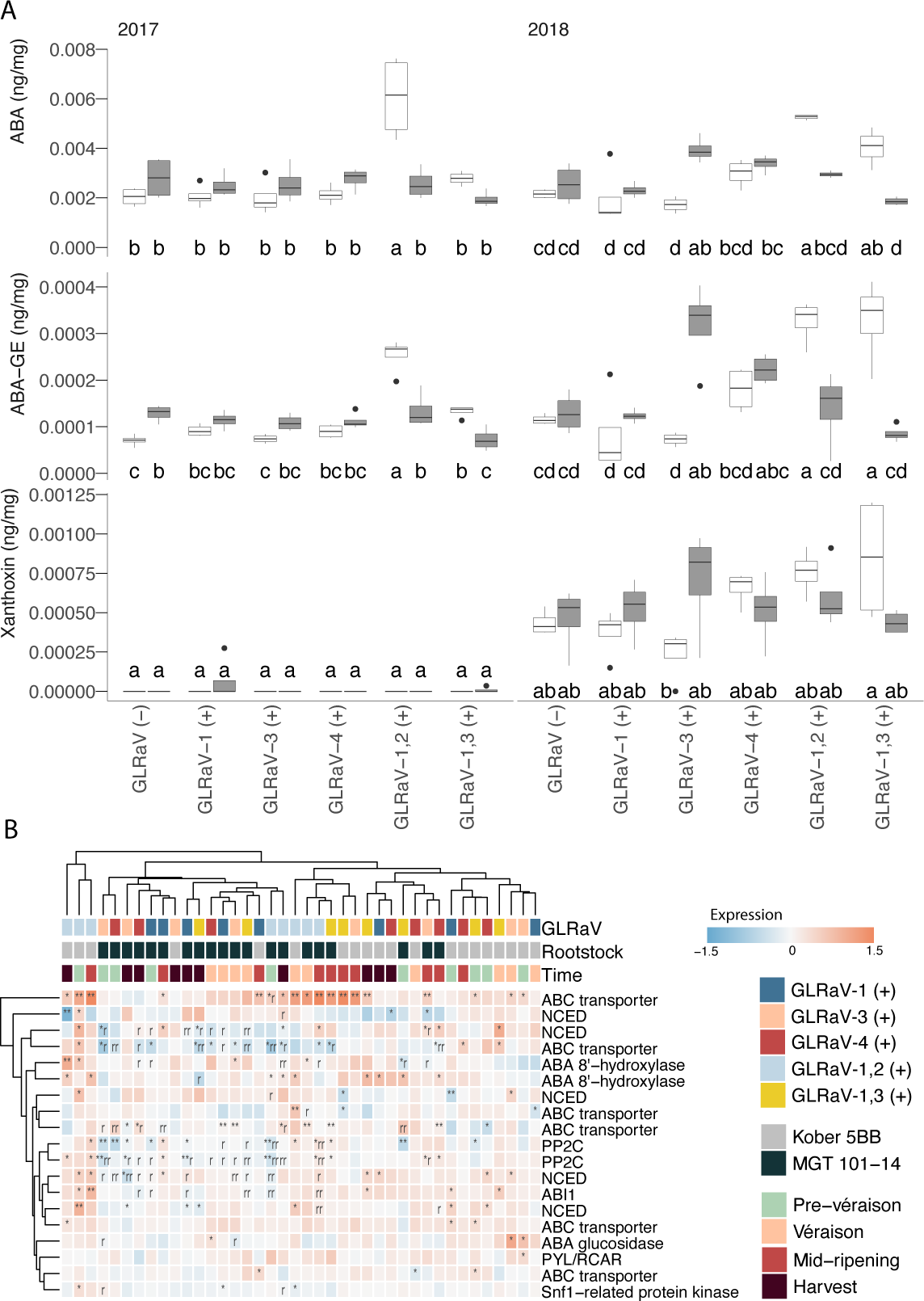
**(A)** Abundance of ABA, ABA-GE, and xanthoxin in 2017 and 2018 at pre-véraison. Groups with non-overlapping letters are significantly different (Tukey HSD, P < 0.05). **(B)** The effect of GLRaV infection(s) and rootstock on ABA biosynthetic and signaling genes. One asterisk, differentially expressed in one year; Two asterisks, differentially expressed in both years. One or two “r”, the effects of a particular GLRaV infection differs between rootstocks at the same developmental stage. Any notation requires the direction (up/down-regulation) of the effect to be consistent in both years, even if a significant change occurred in only a single year.

In the course of development, significant differences in the abundance of these metabolites between rootstock conditions were observed most at pre-véraison and in GLRaV (-), GLRaV-3 (+), and dual infections (Fig. 4A, Supplementary File S8). In GLRaV (-) and most single infection conditions, the level of all three metabolites tended to be higher in berries from plants grafted to MGT 101-14 than in berries from plants grafted to Kober 5BB. The opposite tended to be true when two GLRaVs were present. With one exception, significant changes in the abundance of ABA and ABA-GE in GLRaV (+) versus GLRaV (-) were typically increases and most often observed in berries from Kober 5BB-grafted plants (Tukey HSD, P < 0.05). Though slight reductions in these metabolites were often observed versus GLRaV (-), these were almost always non-significant.

We further investigated abscisic acid biosynthesis and signaling using a previously published, curated set of genes (Pilati *et al*., 2017). Their expression largely clustered according to rootstock, with some exceptions; Kober 5BB GLRaV-1,2 (+), for example, tended to cluster separately from other Kober 5BB-grafted plants. Nonetheless, the effects of GLRaVs differed between rootstocks for many of these genes (Fig. 4B). One of these, an ABC transporter (VITVvi_vCabFran04_v1_P438.ver1.0.g381930), is also included in Figure 3; significant increases in its expression were observed in both years, in both rootstock conditions, and for several GLRaV infections. All other significant changes in GLRaV (+) vs. GLRaV (-) that were reproduced in both years occurred in only one rootstock or the other. These changes were sparse. However, significant differences between rootstocks in identical GLRaV (+) were reproduced in both years for 9 out of these 19 genes. Significant differences between rootstocks in at least one year were observed for 16 out of these 19 genes. On average, three of the 9-*cis*-epoxycarotenoid dioxygenases (NCEDs), both ABA 8’ hydroxylases, three ABC transporter genes, one gene encoding PP2C, and PYL/RCAR were up-regulated in berries from plants infected with GLRaV in both rootstock conditions. One ABC transporter gene was down-regulated in both rootstock conditions. Of the remaining eight genes, most were down-regulated across development only in berries from MGT 101-14-grafted plants.

### Gene expression, hormone, and other metabolite data distinguish the effects of GLRaVs in different rootstocks

Differential expression analysis identified 1,809 genes that (1) were differentially expressed (P < 0.05) in at least one year, (2) in only one rootstock and more than one GLRaV infection type versus GLRaV (-) and/or (3) the effects of more than one GLRaV infection significantly differed between rootstocks (Supplementary Fig. S9). RNAseq, hormone, and metabolite data from ripening Cabernet franc berries were integrated in a multiple factor analysis (MFA) to relate these variables and distinguish the effects of GLRaVs given different rootstocks (Fig. 5, Supplementary Fig. S10, Supplementary Table S11).

**Figure 5.**
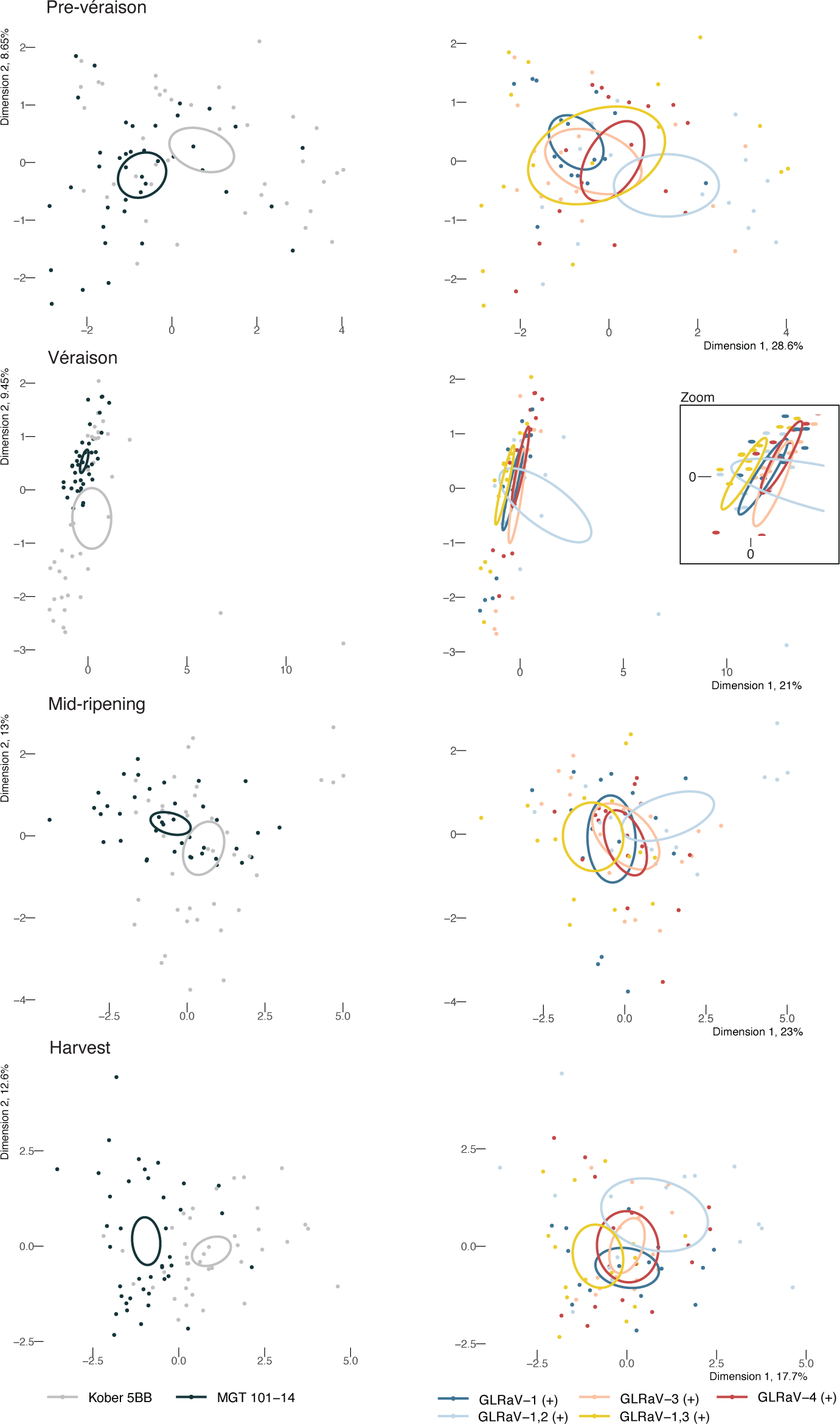
MFA of GLRaV (+) effects. Sample distribution along first two dimensions at each developmental stage. 95% Confidence ellipses are drawn for rootstocks (left) and GLRaV (+) conditions (right).

As input for the MFA, all genes differentially expressed between GLRaV (-) and GLRaV (+) or between rootstocks were used, plus all hormones and metabolites measured. Overall, the rootstocks were distinct at each developmental stage (Fig. 5). Some of the GLRaV (+) conditions could be distinguished from others at pre-véraison, véraison, and harvest. At pre-véraison, GLRaV-1,2 (+) differed overall from GLRaV-1 (+). At véraison, GLRaV-1,3 (+) differed from every other GLRaV (+) condition except GLRaV-1,2 (+). At harvest, the two dual infections were different than one another and GLRaV-1,2 (+) differed from GLRaV-1 (+) (Fig. 5).

Next, we identified which variables were best correlated with each rootstock-differentiating MFA dimension. At each developmental stage, ABA and/or ABA-related metabolites were correlated with at least one of the first two MFA dimensions (Fig. 6A). The rootstock-dependent disparity in ABA and ABA-related genes (Fig. 4) is consistent with the observation that ABA and related metabolites tended to be highly correlated with rootstock-differentiating MFA dimensions over time and that ripening initiates earliest in Kober 5BB plants with dual infections in terms of TSS (Supplementary Fig. S3).

**Figure 6.**
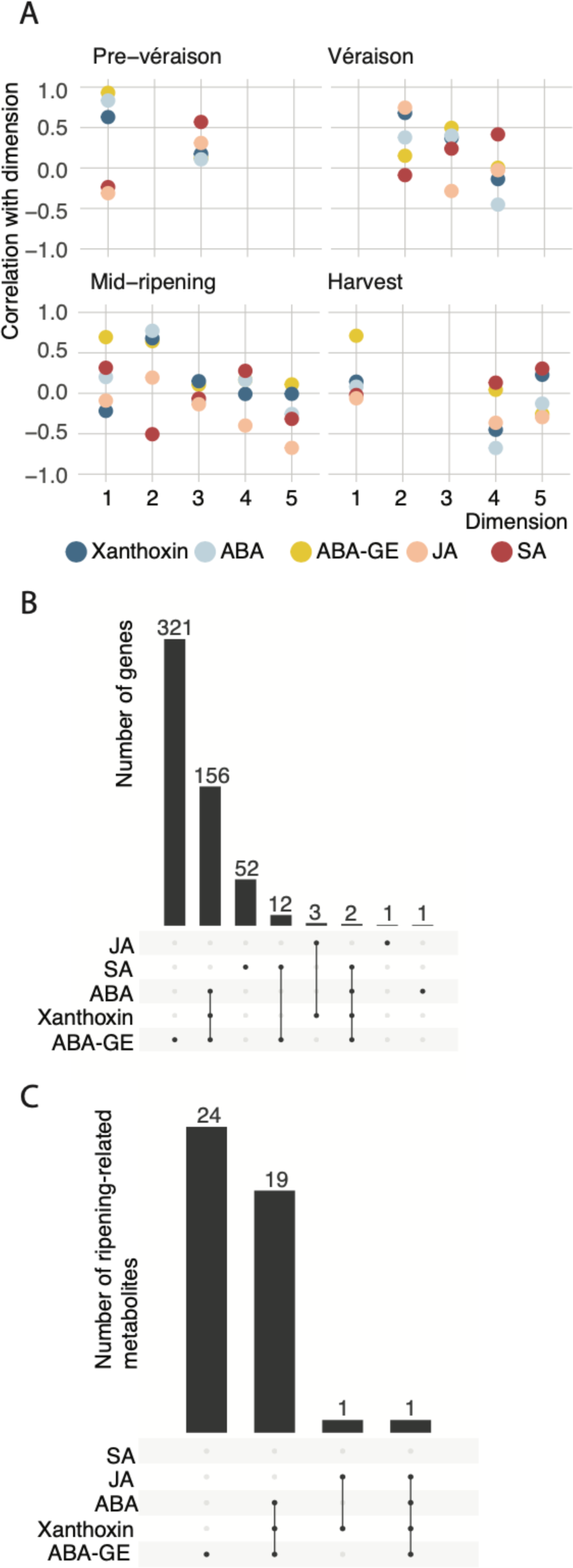
The role of genes, hormones, and metabolites in MFA of GLRaV effects. **(A)** Correlation between hormones and closely related metabolites to rootstock-differentiating MFA dimensions. **(B)** Relationship between genes and hormones, and (C) metabolites and hormones along rootstock-differentiating MFA dimensions. Shown are counts of genes and metabolites correlated to the same dimensions as individual or groups of hormones or hormone-related metabolites.

There were 548 genes that shared high correlation (|corr| > 0.5) to rootstock-differentiating MFA dimensions with hormones or hormone-related metabolites. Most of these genes had shared positive or negative correlations to the same dimensions as ABA, xanthoxin, and/or ABA-GE (Fig. 6B). Genes with functionally relevant relationships to each hormone or hormone-related metabolite were overrepresented (Hypergeometric test, P < 0.05 and n > 1 gene) among the genes that shared high correlation to rootstock-differentiating MFA dimensions with each hormone. Genes with functional categories related to ABA signaling, starch biosynthesis catabolism, and in the C2C2-DOF transcription factor family were overrepresented among the genes correlated to the same dimensions as ABA and xanthoxin. The latter two categories were significantly overrepresented among the genes correlated with the same dimensions as ABA-GE. Genes related to heat shock protein (HSP) mediated protein folding, chaperone-mediated protein folding, cation channel-forming Heat Shock Protein-70, channels and pores, the reductive carboxylate cycle, and carbon fixation were overrepresented among those correlated with the same MFA dimensions as SA. Similarly, the abundance of most ripening-related metabolites measured were correlated with the same MFA dimensions as ABA, xanthoxin, and/or ABA-GE (Fig. 6C).

Overall, the effects of GLRaVs differ between rootstocks primarily in terms of ABA and related metabolites. This finding is especially salient because of the role that ABA plays as a ripening promoter near véraison, in root-scion communication, and plant stress. ABA, metabolites, and genes that were well-correlated to rootstock-differentiating MFA dimensions and were differentially expressed were scrutinized more closely.

### Rootstock influences the impact of GLRaV on hormone signaling genes and transcriptional controls

There were 548 genes in 85 functional categories that were well-correlated with rootstock-differentiating MFA dimensions and differentially expressed between GLRaV (+) grafted to different rootstocks or in GLRaV (+) versus GLRaV (-) in only one rootstock condition (Fig. 7). These functional categories were generally related to hormone and other types of signaling, amino acid and other metabolism pathways, transcription factors, transport, and cellular organization and biogenesis (Fig. 7). Most of these genes coincided with ABA and related metabolites along rootstock-differentiating MFA dimensions (Fig. 7).

**Figure 7.**
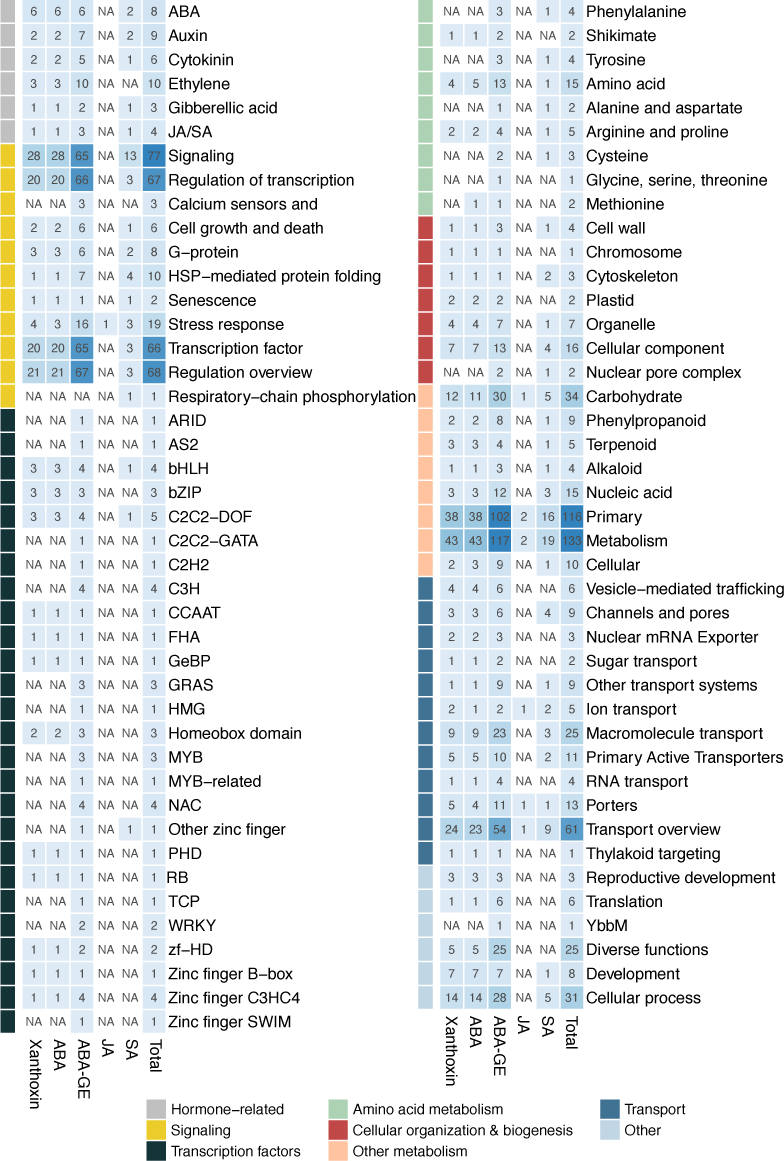
Functional categories of genes correlated to the same rootstock-differentiating MFA dimensions as hormones and hormone-related metabolites. The counts of genes related to each hormone (shared high positive or negative correlation to dimension, (|corr| > 0.5) in each category are shown.

The distribution of expression for four transcription factor families differed significantly between rootstocks at all four developmental stages (Supplementary Fig. S12, Kolmogorov Smirnov, P < 0.05). This included bHLH, C2C2-DOF, FHA, and homeobox domain transcription factors. The distribution of expression for 39 hormone signaling-related genes differed significantly between rootstocks (Fig. 7, Supplementary Fig. S12, Supplementary File S13, Kolmogorov Smirnov, P < 0.05). This was true at each developmental stage for ABA, gibberellin (GA), and auxin signaling genes and at three developmental stages for JA/SA, cytokinin (CK), and ethylene signaling genes (Supplementary Fig. S12). Genes related to all of these hormone families had similar roles in MFA and were associated with ABA, including the JA/SA signaling genes (Fig. 7). This may reflect interactions between hormone signaling pathways. In addition, the effects of GLRaV infections (dual infections and GLRaV-4 (+)) on histone H1 expression were not equal in plants grafted to both rootstocks (Supplementary Fig. S14). Linker histone H1 contributes to higher-order chromatin structure (Hill, 2001).

There were seven ABA signaling pathway genes (excluding VITVvi_vCabFran04_v1_P495.ver1.0.g468110, which was also associated with GA signaling) that differentiated GLRaV effects in plants grafted to different rootstocks (Fig. 8). The mean level of expression for all of these was higher in Kober 5BB than MGT 101-14 (Supplementary Fig. S12). This included *SOS2*, *KEG*, three *PP2C* (*HAB1*, *AHG3*, *DBP*) genes, and genes encoding two ABA-responsive element-binding proteins (*AREB2*, *ABI5*). SOS2 is a kinase appreciated for its role in salt stress response, seed germination, GA signaling (Trupkin *et al*.) and in ABA signal transduction via its interaction with ABI2 and ABI5 (Ji *et al*., 2013; Zhou *et al*., 2015). SOS2 was up-regulated in both rootstock conditions before and at véraison and down-regulated after véraison. KEG is a negative regulator of ABA signaling; it maintains low levels of ABI5 in the absence of stress by ubiquitination and degradation (Stone *et al*., 2006; Lyzenga *et al*., 2013) and helps regulate endocytic trafficking and the formation of signaling complexes on vesicles during stress (Gu & Innes, 2011). KEG was down-regulated in Kober 5BB and down-regulated in MGT 101-14 at véraison and mid-ripening.

**Figure 8.**
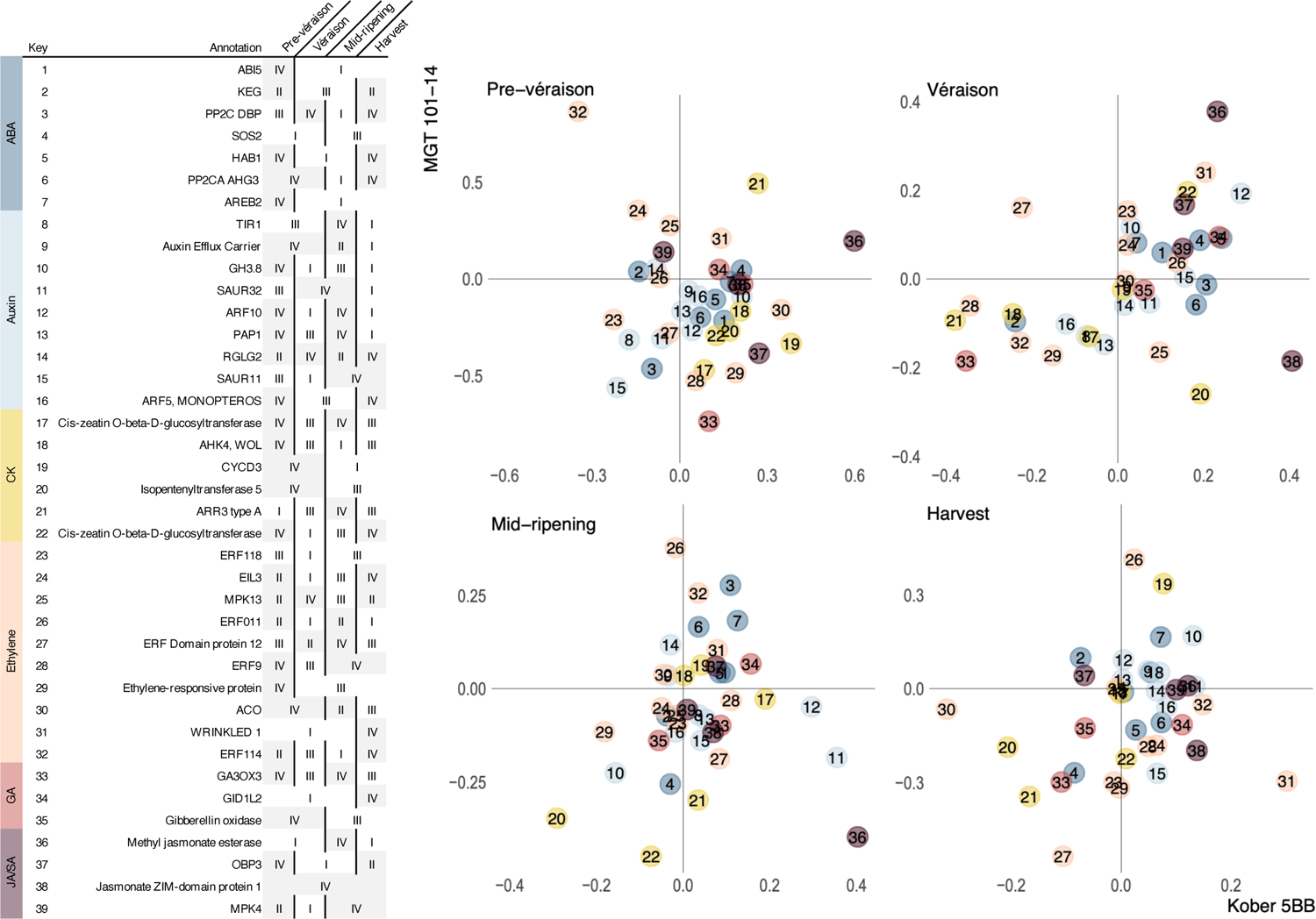
The effect of GLRaVs on hormone signaling gene expression in grape berries from plants grafted to different rootstocks. Quadrants are numbered counterclockwise from top right (I) to bottom right (IV). Individual genes are numbered 1-39.

In the presence of ABA, ABA receptors (PYR/PYL/RCAR family proteins) bind PP2Cs like HAB1 and AHG3 to inhibit their phosphatase activity. As a result, ABA signal transduction is permitted via SnRK2 phosphorylation of ABA-responsive element-binding proteins (Hirayama & Shinozaki, 2010). ABI5 and AREB2 are bZIP transcription factors that bind to ABA responsive elements (ABREs) to drive ABA signaling and ABI5 can integrate signals across hormone signaling pathways (Skubacz *et al*., 2016). The effects of GLRaVs on these genes in Kober 5BB were consistent with an enhancement of ABA signaling during ripening. In Kober 5BB, HAB1 and AHG3 were down-regulated, and AREB2 and ABI5 were up-regulated. In MGT 101-14, the PP2Cs were down-regulated for at least two developmental stages; AREB2 and ABI5 were up-regulated at and after véraison.

### Rootstock influences the impact of GLRaV on the flavonoid biosynthetic pathway

In addition to analyzing hormone and hormone-related metabolites, we analyzed metabolites associated with the shikimate, phenylpropanoid and flavonoid pathways (Fig. 9AB) and their biosynthetic and regulatory genes (Fig. 9C) in Cabernet franc berries during ripening (Supplementary Fig. S15). Significant differences in expression versus GLRaV (-) were detected, as well as significant differences in the effects of GLRaV infection between different rootstock conditions (Fig. 9C). The effects of GLRaV infection on the genes associated with this pathway were generally consistent with the change in abundance of corresponding metabolites (Fig. 9B). Overall, these genes tended to be up-regulated in GLRaV (+) at véraison (Fig. 9C). After véraison, the amount of up-regulation tended to decrease, or genes were down-regulated (Fig. 9C). The three amino acids examined (phenylalanine, tryptophan, and tyrosine) tended to be less abundant in GLRaV (+) across the developmental stages and the largest decreases were observed at harvest (Fig. 9B). Mixed effects of GLRaVs were observed on the abundance of hydroxycinnamic acids (caftaric and coutaric acid), *t-*resveratrol, and anthocyanins. Significant changes versus GLRaV (-) tended to occur in only one year. During ripening, these were significantly more abundant in Kober 5BB GLRaV-1,2 (+), Kober 5BB GLRaV-5 (+), and/or MGT 101-13 GLRaV-3 (+) (Fig. 9B). Significant decreases were observed for GLRaV-1 (+) and GLRaV-1,3 (+). Though non-significant, the size of the downward effect of some GLRaV infections on these metabolites was tended to increase towards harvest. Finally, flavanols (epigallocatechin and catechin) and flavonol (quercetin) glycosides tended to be elevated in GLRaV (+) (Fig. 9B). The size of this effect tended to be greatest before and at véraison and decreased towards harvest. Significant differences between rootstocks were observed for GLRaV-1,2 (+) in both years and for GLRaV-1 (+), GLRaV-1,3 (+), and GLRaV-3 (+) in individual years; the increase in flavonols and flavanols tended to be greater in berries from Kober 5BB GLRaV (+) than MGT 101-14 GLRaV (+).

**Figure 9.**
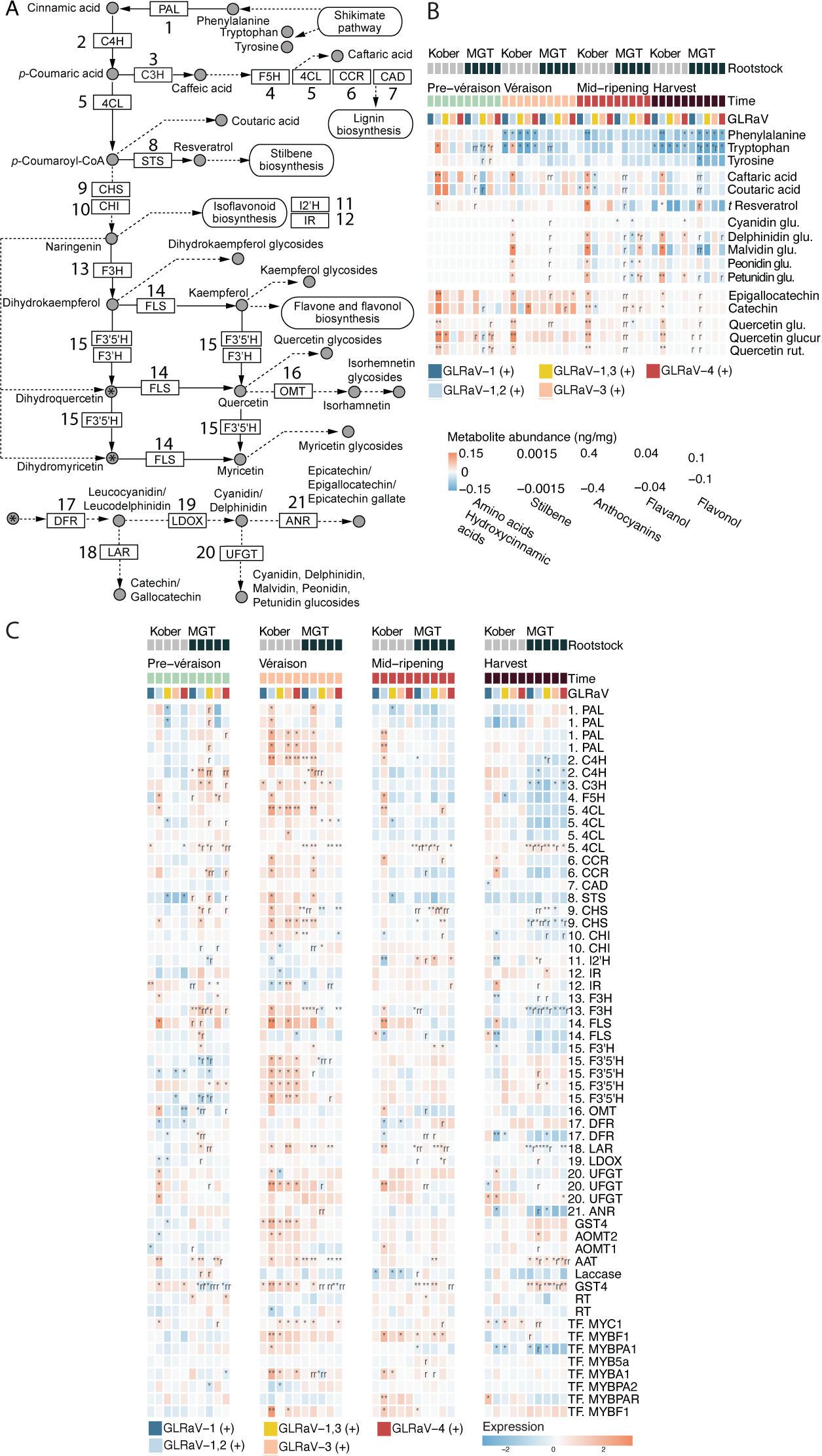
Differentially expressed genes and selected metabolites products of the shikimate, phenylpropanoid, and flavonoid pathways. **(A)** Pathway diagram. **(B)** Metabolite abundances and **(C)** related biosynthetic and regulatory gene expression relative to GLRaV (-) in identical rootstock and at same developmental stage. Notation requires the direction (up/down-regulation) of the effect to be consistent in both years, even if a significant change occurred in only a single year. Glycosides are abbreviated: glu, 3-O-glucoside; glucur, 3-O-glucuronide; rut, 3-O-rutinoside. One asterisk, differentially expressed/abundant in one year; Two asterisks, differentially expressed/abundant in both years. One or two “r”, the effects of a particular GLRaV infection differs between rootstocks at the same developmental stage. PAL, phenylalanine ammonia-lyase; C4H, trans-cinnamate 4-monooxygenase; C3H, p-coumarate 3-hydroxylase; F5H, ferulate-5-hydroxylase; 4CL, 4-coumaroyl-CoA ligase; CCR, cinnamoyl-CoA reductase; CAD, Cinnamyl alcohol dehydrogenase; STS, stilbene synthase; CHS, chalcone synthase; CHI, chalcone isomerase; I2’H, Isoflavone 2’-hydroxylase; IR, Isoflavone reductase; F3H, flavonone 3-hydroxylase; FLS, flavonol synthase; F3’H, flavonoid 3’-monooxygenase; F3’5’H, flavonoid 3’,5’-hydroxylase; OMT, O-methyltransferase; DFR, dihydroflavanol 4-reductase; LAR, leucoanthocyanidin reductase; LDOX, Leucoanthocyanidin dioxgenase; UFGT, UDP-glucose:anthocyanidin/flavonoid 3-O-glucosyltransferase; ANR, anthocyanidin reductase; GST4, Glutathione S-transferase 4; AOMT, Anthocyanin O-methyltransferase; AAT, Anthocyanin acyl-transferase; RT, UDP-rhamnose: rhamnosyltransferase. The pathway annotation is based on KEGG pathways (www.genome.jp/kegg/pathway.html, accessed February 13, 2021) and Blanco-Ulate *et al.,* 2017.

## Discussion

Grapevine leafroll-associated viruses affect viticulture on nearly every continent (Habili & Nutter, 1997; Akbaş *et al*., 2007; Fiore *et al*., 2008; Mahfoudhi *et al*., 2008; Golino *et al*., 2008a; Charles *et al*., 2009; Jooste *et al*., 2015) and can cause considerable economic impact on a major crop.

The presence and severity of symptoms in GLRaV-infected grapevines is influenced by host genotype, rootstock, which GLRaV is present, and environmental conditions. In addition to the assembly and annotation of the Cabernet franc genome, a valuable resource that might be applied for the larger purpose of understanding grapevine genomic diversity and evolution, the dedicated experimental vineyard used in this study is a tremendous asset for the study of GLRaV infections over time in a common environment. This work identified responses to GLRaVs in grape berries during ripening, including those that are conserved given different GLRaVs and rootstocks and responses that differ based on the rootstock present. We propose which hormones and signaling pathways at least partially govern the responses observed and likely influence leafroll disease symptoms.

The effects of dual infections, particularly GLRaV-1,2 (+), were most distinctive. All of the leafroll viruses selected for this study belong to the *Closteroviridae* family and all but one belongs to the *Ampelovirus* genus; GLRaV-2 belongs to *Closterovirus*. All of the GLRaVs used in this study contain a relatively consistent replication gene block (RGB) but are markedly diverse outside of the RGB (Naidu *et al*., 2015). In addition to host genotype and environment, the sequences downstream of the RGB may account for the disparities in responses observed between infection types. These sequences encode a quintuple gene block and/or viral suppressors of host RNA silencing (VSRs; Naidu *et al.,* 2015). Some of are characterized (Gouveia *et al*.), but most are not. Further research might determine their specific effects and relationship to host cellular machinery (Chapman, 2004; Wu *et al*., 2010).

Changes in the expression of NBS-LRR genes were among the conserved responses to GLRaVs and were the single largest category of genes among them. NBS-LRR genes confer resistance to powdery and downy mildew in grapevine (Riaz *et al*., 2011; Zini *et al*., 2019). The abundance of these genes varies among *Vitis* species and are particularly dense at resistance loci (Cochetel *et al*., 2021). Hypersensitive response (HR) to viruses is mediated by R genes. SA subsequently accumulates and pathogenesis-related gene expression increases for systemic acquired resistance. HR is a means of prohibiting pathogen spread and can confer resistance when a corresponding dominant avirulence protein is produced by the pathogen (Moffett *et al*., 2002; Balint Kurti, 2019). However, GLRaV infections are systemic, persist over time, and SA does not seem to play a preeminent role in the response to GLRaV infections. In a previous study of GLRaV-3 infections in Cabernet Sauvignon and Carmenère, the authors also remarked on the induction of defense genes but their inability to impede systemic infection (Espinoza *et al*., 2007b). Both SA and ABA can participate in virus response, though considerably less is understood about the role of ABA (Kunkel & Brooks, 2002; Koornneef & Pieterse, 2008; Alazem & Lin, 2015) and its relationship to NBS-LRRs, specifically. Notably, however, ABA deficiency is associated with an increase in R-gene efficacy in incompatible interactions with *Pseudomonas syringae* and in a manner independent of SA (Mang *et al*., 2012).

Hormones have been implicated in mediating defense and development-related networks and are overrepresented at network hubs (Amrine *et al*., 2015; Müller & Munné-Bosch, 2015; Jiang *et al*., 2015; Vandereyken *et al*., 2018). Hormones like ABA, SA, JA, and others act as important signaling molecules during ripening and defense. The pathways engaged under stress are often tailored to particular pathogens and this entails coordination between hormone pathways (Gao *et al*., 2011; Vos *et al*., 2015). Interestingly, the effects of GLRaVs on gene expression in several hormone signaling pathways differed between rootstocks. A subsequent effort could be made to measure the abundances of hormones not quantified here, like cytokinin, gibberellin, and ethylene. Of the hormones considered in this study, however, ABA and ABA-GE abundance tended to increase in GLRaV (+) and was influenced by rootstock. ABA can antagonize SA and JA signaling pathways and suppress ROS signaling (Alazem & Lin, 2015). WRKY transcription factors both regulate and/or are regulated by ABA, SA, and JA (Li, 2004; Jiang & Deyholos, 2008; Gao *et al*., 2011; Liu *et al*., 2015; Xin *et al*., 2016). SA and ABA both can interact with RNAi, which is a fundamental component of anti-viral defense. *AGO1* expression is positively correlated with ABA and *miR168a*, which regulates AGO1, contains ABA-responsive regulatory elements in its promoter (Alazem & Lin, 2015). Levels of ABA and ABA-GE increase in tobacco mosaic virus-infected leaves. One way in which ABA might aid plant defense is by increasing callose deposition to impair virus movement (Alazem & Lin, 2015); a gene encoding callose synthase is upregulated in the leaves of grapevine virus B-infected plants (Chitarra *et al*., 2018).

Relatively more is known about ABA’s function as a ripening promoter (Wheeler *et al*., 2009; Koyama *et al*., 2010), in response to drought stress (Deluc *et al*., 2009a; Cochetel *et al*., 2020), and in transmitting long-distance signals from roots to aerial organs and vice versa (Manzi *et al*., 2015; Ferrandino *et al*.). In a study of the impact of GLRaV-3 infection, drought stress, and a combination of both on *in-vitro* grapevine plantlets, individual stresses both induced ABA (Cui *et al*., 2016). Drought stress increases ABA and induces the flavonoid pathway in both tea plants (Gai *et al*., 2020) and in grapes (Deluc *et al*., 2009b). Our findings, in which ABA abundance tends to increase in GLRaV (+), are different than that observed for Red Blotch virus-infected berries, in which ABA abundance and *NCED* expression decrease in infected fruits (Blanco-Ulate *et al*., 2017).

The results of our analysis of metabolites associated with the phenylpropanoid and flavonoid pathways are mixed in their consistency with previous work. Though nonsignificant decreases in anthocyanins were observed, anthocyanins significantly increased in several GLRaV (+) conditions, albeit usually in individual years. These findings differed from others; some observed significant decreases in anthocyanins in fruits from GLRaV-infected plants (Lee & Martin, 2009; Vega *et al*., 2011) and others observed no significant changes in anthocyanin at harvest (Endeshaw *et al*., 2014; Alabi *et al*., 2016). Like Vega *et al.,* (2011), flavonols were elevated in GLRaV (+) and the largest differences versus GLRaV (-) occurred at the first two stages, *FLS* expression was down-regulated in GLRaV (+) at harvest, and *CHS* and *MYBPA1* expression were up-regulated at véraison and generally down-regulated at harvest.

In this study, changes in the abundance of ABA and related metabolites distinguished the effects of GLRaVs between rootstocks. The parentage of the two rootstocks used in this study, Kober 5BB and MGT 101-14, includes *V. riparia.* The other parents of Kober 5BB and MGT 101-14 are *V. berlandieri* and *V. rupestris,* respectively. These rootstocks were developed at different times. MGT 101-14 originated in France in 1882 and Kober 5BB originated in Austria in 1930 (https://fps.ucdavis.edu/). Rootstocks are chosen for the advantages they confer to the scion given a particular set of circumstances, often having to do with resistance to *Phylloxera*, nematodes, scion vigor, soil type, and abiotic stress tolerance (Tramontini *et al*., 2013; Corso & Bonghi, 2014; Warschefsky *et al*., 2016). Unlike *V. vinifera*, *V. riparia, V. rupestris, and V. berlandieri* are asymptomatic hosts of GLRaVs (Naidu *et al*., 2014). Yet, the particularly severe response to GLRaV-1,2 (+) was not entirely unexpected in Kober 5BB-grafted vines (Uyemoto *et al*., 2001; Golino *et al*., 2008b; Alkowni *et al*., 2011). Differences in wood abnormalities given different rootstocks (including Kober 5BB) have been observed for particular isolates causing grapevine rugose wood disease (Credi, 1997). Notably, nutritional deficiencies in phosphorus, magnesium, and potassium produce symptoms that resemble those typically observed in GLRaV-infected plants (Gohil *et al.,* 2016). Magnesium deficiency tolerance (Livigni *et al*., 2019), the impact of phosphorus deficiency on canopy growth (Grant & Matthews, 1996), and potassium uptake and channels are all influenced by rootstock (Wolpert *et al*., 2005). Potassium uptake, channel activity, and related gene expression are also regulated by ABA (Blatt, 2000; Köhler *et al*., 2003; Song *et al*., 2016; Rogiers *et al*., 2017), and the application of ABA to tomato roots by drip irrigation affects fruit mineral composition (Barickman *et al*., 2019). Furthermore, elevated levels of potassium are observed in leafroll virus-infected Burger and Sultana and Burger fruits (Kliewer & Lider, 1976b; Hale & Woodham, 1979) and in leaf petioles but lower in leaf blades (Cook and Goheen, 1961). Perhaps potassium-deficient and GLRaV (+) phenotypes are similar because they both induce a response governed by ABA that can be fine-tuned by rootstocks. If some portion of scion ABA originates in roots and/or if rootstock can influence scion ABA levels and signaling genes, as observed here and by others (Chitarra *et al*., 2017). then perhaps this partially accounts for the variation in response observed between rootstocks. This experiment did not include a comprehensive survey of phytohormones, which would be beneficial, but ABA’s function in “root-shoot” communication, its role in ripening, and the results here make it a good candidate around which to study the basis of leafroll disease symptom variability going forward. In addition, the transport of RNAs across the graft junction may perform some function that affects scion disease severity, but this remains to be seen in the particular case of GLRaV (Chitarra *et al*., 2017).

Together, these data support several conclusions. (1) The majority of genes differentially expressed as a consequence of infection or between GLRaV (+) plants with different rootstocks were year-specific. A small subset of effects was consistently observed across experimental conditions and in both years. These shared changes in expression involved genes associated with pathogen detection, ABA signaling and transport, ROS-related signaling, cytoskeleton remodeling, vesicle trafficking, phenylpropanoid metabolism, sugar transport and conjugation, and leucine biosynthetic genes. (2) The impacts of GLRaV-1,2 dual infection on Kober 5BB-grafted vines were the most distinctive and severe. Though there was variation between GLRaV infections observed, only the effects of GLRaV-1,2 were distinguishable overall from other infections (GLRaV-1 and/or GLRaV-1,3) pre-véraison and at harvest. (3) The particular effects of GLRaVs in plants grafted to different rootstocks were distinguishable overall at every developmental stage. ABA-related variables were among those that best distinguished the responses to GLRaVs in different rootstock conditions. This included the abundance of ABA, the abundance of ABA-GE, and the expression of genes associated with ABA and other hormone signaling pathways. Finally, this work alone is insufficient to recommend the use of one rootstock or another, but the disparity in sensitivity and symptom severity observed in berries from Cabernet franc vines grafted to different rootstocks suggests that rootstock selection generally should be further explored as a strategy to mitigate some of the negative consequences of leafroll virus infections, should vectors of the virus encroach upon a vineyard.

## Supplementary Data

*Supplementary File S1*. Curated list of ABA-related genes. Pilati et al., 2017

*Supplementary File S2.* Hormone and metabolite retention time and mass information.

*Supplementary Fig. S3*. Malic acid, moisture content, pH, titratable acidity, weight per berry, yeast assimilable nitrogen (YAN), and total anthocyanins in 2015 and 2016. For most measures in both years, 3<= n <=9. Total anthocyanin and moisture content in 2015, 2<=n<=9. Total soluble solids before harvest in 2017 and 2018, n=6. Groups differences are indicated with non-overlapping letters (Tukey HSD, P < 0.05).

*Supplementary Table S4*. Cabernet franc assembly and annotation statistics.

*Supplementary Fig. S5*. Pearson correlation between RNAseq samples

*Supplementary Fig. S6*. Barplots showing the number of differentially expressed (P < 0.05) genes up and down-regulated in MGT 101-14 grafted plants versus Kober 5BB grafted plants for each GLRaV condition in each year and in both years.

*Supplementary File S7.* Conserved responses to GLRaVs depicted in Fig. 3B.

*Supplementary File S8.* Abundance of ABA, ABA-GE, and xanthoxin in 2017 and 2018 at véraison, mid-ripening, and at harvest. Abundance of ABA, ABA-GE, and xanthoxin in 2017 and 2018 at all four developmental stages. ANOVA tables for all 5 hormones and hormone-related metabolites. Groups with non-overlapping letters are significantly differet (Tukey HSD, P < 0.05).

*Supplementary Fig. S9*. Rootstock-differentiating genes

*Supplementary Fig. S10*. Percent of variation explained by each of the first five MFA components at each developmental stage.

*Supplementary Table S11*. Number of MFA variables positively and negatively correlated with each rootstock-differentiating MFA dimension.

*Supplementary Fig. S12*. Distribution of centered expression for genes, per functional category, correlated to the same rootstock-differentiating MFA dimensions as hormones and hormone-related metabolites. Results of Kolmogorov-Smirnov test are shown.

*Supplementary Text S13.* Additional description of hormone signaling genes in Figure 8.

*Supplementary Fig. S14*. Centered expression of VITVvi_vCabFran04_v1_P478.ver1.0.g430590, Histone H1, in berries from GLRaV (+) plants grafted to two different rootstocks.

*Supplementary Fig. S15*. All metabolite products measured that are associated with the shikimate, phenylpropanoid, and flavonoid pathways. One asterisk, differentially expressed/abundant in one year; Two asterisks, differentially expressed/abundant in both years. One or two “r”, the effects of a particular GLRaV infection differs between rootstocks at the same developmental stage.

## Acknowledgements

This work was funded by the Pierce’s Disease and Glassy-winged Sharpshooter Research Board of the California Department of Food and Agriculture (project 17-0417-000-SA). DC was partially supported by the Louis P. Martini Endowment in Viticulture. The authors would like to thank all past and present Cantu lab members who assisted with the laborious process of sampling, deseeding, crushing, and weighing the tissue used in this study, including Abraham Morales-Cruz, Eric Tran, Jerry Lin, Barbara Blanco-Ulate, and Lucero K. Espinoza.

## Author contributions

AR, DAG, SEE, MAR, and DC designed the experiments. MAR monitored virus infections. MM designed the sampling procedure and photographed experimental vines. JG and DQ crushed tissue and prepared RNA extracts. RFB prepared the Cabernet franc genomic and RNA sequencing libraries. AM assembled and annotated the Cabernet franc genome. LL created the metabolite acquisition method. AMV and LL generated the metabolite data. AMV analyzed the RNA sequencing data and prepared all the figures. AMV, LL, and DL analyzed the metabolite data. AMV and DC wrote the manuscript. All authors contributed to, read, and approved the final manuscript.

## Data availability

The DNA (BioProject PRJNA701939) and RNA (BioProject PRJNA701940) sequencing data that support these findings are available at the National Center for Biotechnology Information (NCBI). A Cabernet franc genome browser and on-line BLAST tool are currently available at http://www.grapegenomics.com. While under review, access to the Cabernet franc genome assembly and feature sequences, primary scaffolds, haplotigs, feature annotations are available at http://169.237.73.197/for_reviewers/Vondras_leafroll. The assembly, annotations, and raw RNA sequencing counts per library and per gene are available at Zenodo (http://doi.org/10.5281/zenodo.4555977).

